# Transcriptional response of human articular chondrocytes treated with fibronectin fragments: an *in vitro* model of the osteoarthritis phenotype

**DOI:** 10.1101/2020.06.18.155390

**Authors:** Kathleen S. M. Reed, Veronica Ulici, Cheeho Kim, Susan Chubinskaya, Richard F. Loeser, Douglas H. Phanstiel

## Abstract

**Objective:** Fibronectin is a matrix protein that is fragmented during cartilage degradation in osteoarthritis (OA). Treatment of chondrocytes with fibronectin fragments (FN-f) has been used to model OA *in vitro*, but the system has not been fully characterized. This study sought to define the transcriptional response of chondrocytes to FN-f, and directly compare it to responses traditionally observed in OA.

**Design:** Normal human femoral chondrocytes isolated from tissue donors were treated with either FN-f or PBS (control) for 3, 6, or 18 hours. RNA-seq libraries were compared between time-matched FN-f and control samples in order to identify changes in gene expression over time. Differentially expressed genes were compared to a published OA gene set and used for pathway, transcription factor motif, and kinome analysis.

**Results:** FN-f treatment resulted in 1,224 differentially expressed genes over the time course. Genes that are up- or downregulated in OA were significantly up- (p < 0.00001) or downregulated (p < 0.0004) in response to FN-f. Early response genes were involved in proinflammatory pathways and their promoters were enriched for NF-κB-related motifs, whereas many late response genes were involved in ferroptosis, and their promoters were enriched for Jun-related motifs. Highly upregulated kinases included CAMK1G, IRAK2, and the uncharacterized kinase DYRK3, while growth factor receptors TGFBR2 and FGFR2 were downregulated.

**Conclusions:** FN-f treatment of normal human articular chondrocytes recapitulated many key aspects of the OA chondrocyte phenotype. This *in vitro* model is promising for future OA studies, especially considering its compatibility with genomics and genome-editing techniques.

## Introduction

Osteoarthritis (OA) is the most common form of joint disease and affects over 250 million people worldwide, including over 10% of those older than 60 years ^1^. There is no known cure, and treatments are currently limited to symptom management. One major reason for the lack of treatments is an incomplete understanding of the mechanisms that promote OA and its progression. While mouse models and human samples have provided valuable insights into OA biology, new human disease models amenable to manipulation and high-throughput screening would improve our ability to understand and potentially better treat this painful and disabling disease.

OA involves many, if not all, of the tissues that comprise articular joints, with the degradation and loss of articular cartilage noted as a central feature ^2^. Studies of potential OA pathways often compare chondrocytes isolated from normal cartilage obtained from various animal species, including humans, to chondrocytes obtained from OA tissue. A limitation, particularly with human tissue, is that the OA chondrocytes are most often isolated from cartilage obtained at the time of joint replacement, resulting in comparisons being made to cells at an advanced stage of disease. Animal models, including mice, have been critical for mechanistic studies, but major differences in genomes, body structures, and OA prevalence limit the relevance to human biology^3 3^.

A commonly used option for modeling the chondrocyte OA phenotype has been to stimulate primary cells or cell lines *ex vivo* with cytokines such as IL-1 or TNFα ^4^. A major limitation of these studies is that the cells are treated with levels of cytokines in the ng/ml range to obtain a desired response, while (at least in the synovial fluid) IL-1 and TNFα are only present in pg/ml amounts ^5^. In addition, recent studies, including failed clinical trials of IL-1 and TNFα inhibition in OA, suggest that multiple pro-inflammatory mediators contribute to OA development, and IL-1 or TNFα may not be the driving factors ^6–8^.

An alternative *in vitro* model for simulating a chondrocyte OA phenotype utilizes fragments of fibronectin. Fibronectin is an extracellular matrix protein present in cartilage that is upregulated in OA tissue and subsequently degraded by several proteases ^9,10^. Fibronectin fragments (FN-f) of various sizes and at levels in the µM range have been detected in OA cartilage and synovial fluid as well as in cartilage from patients with rheumatoid arthritis^11–13^. Injection of FN-f into rabbit joints was found to induce cartilage proteoglycan loss, which is a feature of early OA^14^. Treatment of isolated human chondrocytes or cartilage explants with FN-f has been shown to recapitulate many known features of OA, including production of multiple matrix-degrading enzymes and proinflammatory cytokines found in OA joints ^9,15,16^. While these results demonstrate the value of FN-f treatment for studying OA, the global similarity between FN-f-treated chondrocytes and OA chondrocytes has not been fully explored.

The purpose of this study was to characterize the transcriptional response to FN-f stimulation of *ex vivo* human chondrocytes and to compare this response to those previously observed in OA. We found that FN-f triggers a robust transcriptional response in primary chondrocytes, which correlates with changes observed during OA. Analysis of gene ontology terms, signaling pathways, and transcription factor motifs revealed that known regulators of OA progression also play a role in the FN-f response, as do a host of genes and pathways that had not previously been implicated in OA. These results support FN-f treatment as a viable model for studying transcriptional control of OA progression and provide a valuable resource for future studies.

## Methods

### Sample collection and treatment

Primary articular chondrocytes were isolated by enzymatic digestion from normal human femoral cartilage obtained from three tissue donors without a history of arthritis and with ages from 50–61 years, as previously described^17^. Cells were cultured to confluency in standard media with 10% fetal bovine serum and then made serum-free for 2 hours prior to treatment with a purified 42 kDa endotoxin-free recombinant FN-f (1 µM in PBS), prepared as previously described, or PBS as a control^18^. The FN-f used here consists of domains 7–10 in native fibronectin which contains the RGD cell-binding domain recognized by the α5β1 integrin. After 3, 6, or 18 hrs of treatment with FN-f or PBS, media was removed, cultures were quickly rinsed with cold PBS, and RNA was immediately isolated using the RNeasy kit from Qiagen.

### RNA-seq library preparation

Prior to library preparation, all RNA samples were analyzed using a Tapestation RNA HS tape to confirm RNA integrity numbers (RIN) were within 8.5–10, which indicates high quality, intact RNA. Ribosomal RNA was removed using the New England Biolabs NEBNext rRNA Depletion Kit (Human/ Mouse/Rat), and RNA-seq libraries were prepared using the NEBNext Ultra II Directional RNA Library Prep Kit and NEBNext Multiplex Oligos for Illumina. Final libraries were then quantified using a Qubit 4 Fluorometer and run on a Tapestation D1000 HS tape, to confirm average fragment sizes were within 260–320 base pairs and calculate molarity for pooling.

### Processing of RNA-Seq libraries

RNA-seq libraries were sequenced to an average depth of approximately 58 million reads per sample (50 base pairs, paired-end reads) on an Illumina HiSeq 4000 (High Output). Low-quality reads and adapters were trimmed using Trim Galore! (v. 0.4.3), and trimmed reads were then quantified using Salmon quasi-mapping (v. 0.8.2). Both programs were run with default settings.

### Identifying differential genes

Gene-level quantifications were summarized from each sample using tximport (v. 1.2.0). Differential analysis was conducted in R with DESeq2 (v. 1.22.2) using a design adjusting for donor variability when calculating differences between treatment groups (∼ donor + treatment). Differential genes were defined as genes with an FDR-adjusted p-value below .01 (Wald test) and an absolute fold-change above 2 when comparing FN-f treated samples to their time-matched controls.

### Temporal clustering of genes

To assign temporal response classes for the 1,224 differential genes, first a z-score was calculated from the variance-stabilized counts, centering the counts in each sample relative to the average counts among all samples for each gene. Then, for each donor sample, the untreated control score was subtracted from the FN-f treated score for every gene. The difference in z-score was then averaged over the three donors, ultimately providing three values for every gene representing the normalized expression relative to the control at each time point (3, 6, and 18 hours). This matrix was then clustered using k-means clustering with a *k* of 4. These clusters were labeled “Up Early”, “Up Late”, “Down Early”, and “Down Late” based on their expression relative to untreated controls at each time point.

### Comparing with genes differentially expressed in OA cartilage

The previously published RAAK study identified genes that were up- or downregulated in OA-affected cartilage compared to preserved cartilage in the same joint^19^. To focus on only the genes that exhibited the strongest changes in OA, the genes from the RAAK study were filtered to include only those that had a p-value of less than 0.01 and an absolute fold change of greater than 1.5. A Mann-Whitney U test was used to determine if the FN-f-induced fold change of each set of OA-responsive genes were significantly higher or lower, respectively, than the FN-f-induced fold change of genes outside of each gene set.

### GO, KEGG & Transcription Factor Motif Enrichment Analysis

The “findMotifs.pl” tool in the HOMER software suite (v. 4.10.4) was used on each cluster of genes in order to identify enriched Gene Ontology (GO) terms, Kyoto Encyclopedia of Genes and Genomes (KEGG) pathways, and transcription factor motifs. Using the “-mknown” option, known transcription factor motifs were detected from the core mononucleotide human HOCOMOCO dataset (v. 11, p-value < 0.001). Transcription factor families were then identified via TFClass^20^. *De novo* motifs with p-values below 1×10^*-8*^ and best-match motif scores below 0.6 were classified as “unannotated”, in that they did not appear to have a conclusive known motif match.

### Kinome Visualization

To identify and visualize protein kinases present in each cluster, the cluster assignments were plotted using the human kinome visualization tool, Coral^21^. Flat text files were created listing the ENSEMBL ID, k-means cluster, and maximum absolute fold change among all three time points for each differential gene. These lists were then used to plot both categorical (cluster, encoded in branch/node color) and qualitative data (maximum absolute fold change, encoded in node size) on the protein kinase tree, originally published by Manning et al ^22^.

### Data Availability

Raw sequencing data, transcript-level quantification output from Salmon, and a table containing gene-level summaries of read counts in each sample, as well as Wald test p-values, log2 fold change, and normalized count z-score at each time point, are made publicly available at GEO accession GSE150411.

## Results

### FN-f induces global changes in chondrocyte gene expression

To determine the extent to which FN-f treatment alters transcription in human chondrocytes, we performed a three-point RNA-seq time course of 3, 6, and 18 hours (**Fig S1**). We used principal component analysis (PCA) of the 18 samples to determine the extent to which each attribute (donor, time in culture, and time treated with FN-f) contributed to transcriptional state. Untreated samples clustered largely by donor rather than by time in culture, suggesting that time in culture had only minor impacts on gene expression (**Fig 1A**). Conversely, samples treated with FN-f clustered by treatment time, suggesting that FN-f treatment played a larger role in transcriptional state than the genotype of the donor cells. Statistical analysis of differential expression patterns confirmed these results. Untreated samples exhibited only 28 differentially expressed genes between time points (FDR < 0.01, fold change > 2; **Fig 1B**). In contrast, comparison of FN-f-treated samples to their time-matched controls revealed 1,224 genes that changed significantly in response to FN-f treatment in at least one of the three time points (**Table S1**). Increased treatment time correlated with increased numbers of differentially expressed genes, with 317, 747, and 850 genes affected at 3, 6, and 18 hours, respectively. Together, these results demonstrate that FN-f treatment has a profound effect on transcription that is distinct from the effects of *ex vivo* culturing, and this effect is robust when accounting for variation in response among biological replicates.

**Figure 1.**
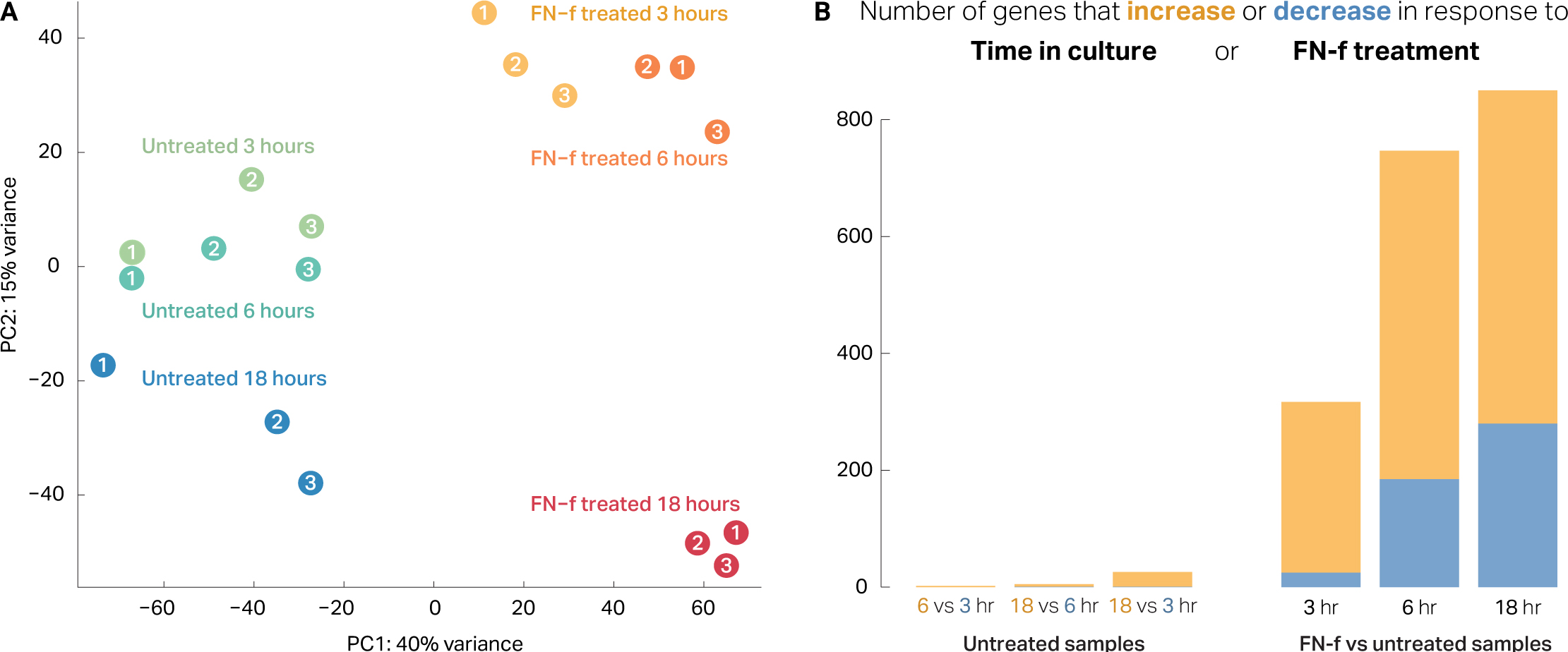
FN-f treatment induces a robust transcriptional response in human chondrocytes. **(A)** Principal component analysis (PCA) of each sample (colors indicate condition, numbers indicate donor) reveals a distinct separation between FN-f-treated and untreated samples along the first principle component (explaining 40% variance). Additionally, FN-f-treated samples cluster by length of treatment rather than by donor, whereas untreated samples cluster more based on donor (particularly along PC1). **(B)** Bar plots depicting the number of genes that exhibit significant differences in expression due to time in culture (left) or FN-f treatment (right; Wald test FDR > 0.01, absolute fold change > 2). Yellow and blue bars represent the number of genes that increase or decrease in each comparison. (Left) A bar plot depicting the number of genes that change significantly between untreated control samples reveals that time in culture has only a minimal impact on gene expression. (Right) A bar plot depicting the number of genes that change significantly in FN-f-treated samples compared to time-matched controls demonstrates that FN-f induces a robust transcriptional response in chondrocytes.

Investigation of specific differential genes revealed expected changes for many known regulators of OA. Genes upregulated in response to FN-f included cytokines and chemokines such as *CXCL2* (189-fold), *LIF* (160-fold), and *IL6* (87-fold). Interleukin-1β (*IL1B*), a proinflammatory cytokine with elevated expression in OA chondrocytes^23,24^, was upregulated at all time points, peaking at over 100-fold. Upregulated genes also included proteases such as *MMP13* and *MMP10* (both 12-fold). Matrix metallopeptidase 13 (*MMP13*) is an enzyme that degrades type II collagen and is thought to play a critical role in cartilage degradation in OA^12,15,25–28^. Interestingly, among the downregulated genes were the collagen-binding integrins *ITGA10* and *ITGA11*, which decreased 3.7- and 3.5-fold, respectively.

### FN-f treatment induces transcriptional changes similar to OA

Changes in gene expression in response to FN-f were compared to the differential expression reported in chondrocytes isolated from OA and preserved tissue in the RAAK study^19^ using a subset of differential genes with the strongest effects (p-value < 0.01, absolute fold change > 1.5) (**Table S2; Fig 2**). Genes upregulated in the OA tissue were also upregulated in response to FN-f treatment (Mann-Whitney U test, p-value < 0.00001), and these effects grew more pronounced with longer exposure to FN-f (**Fig 2**). Genes that were down-regulated in OA tissue were similarly downregulated in response to FN-f, though only exhibiting statistical significance after 18 hours of treatment (Mann-Whitney U test, p-value = 0.0004). These results suggest that FN-f induces similar transcriptional changes to those found in OA chondrocytes.

**Figure 2.**
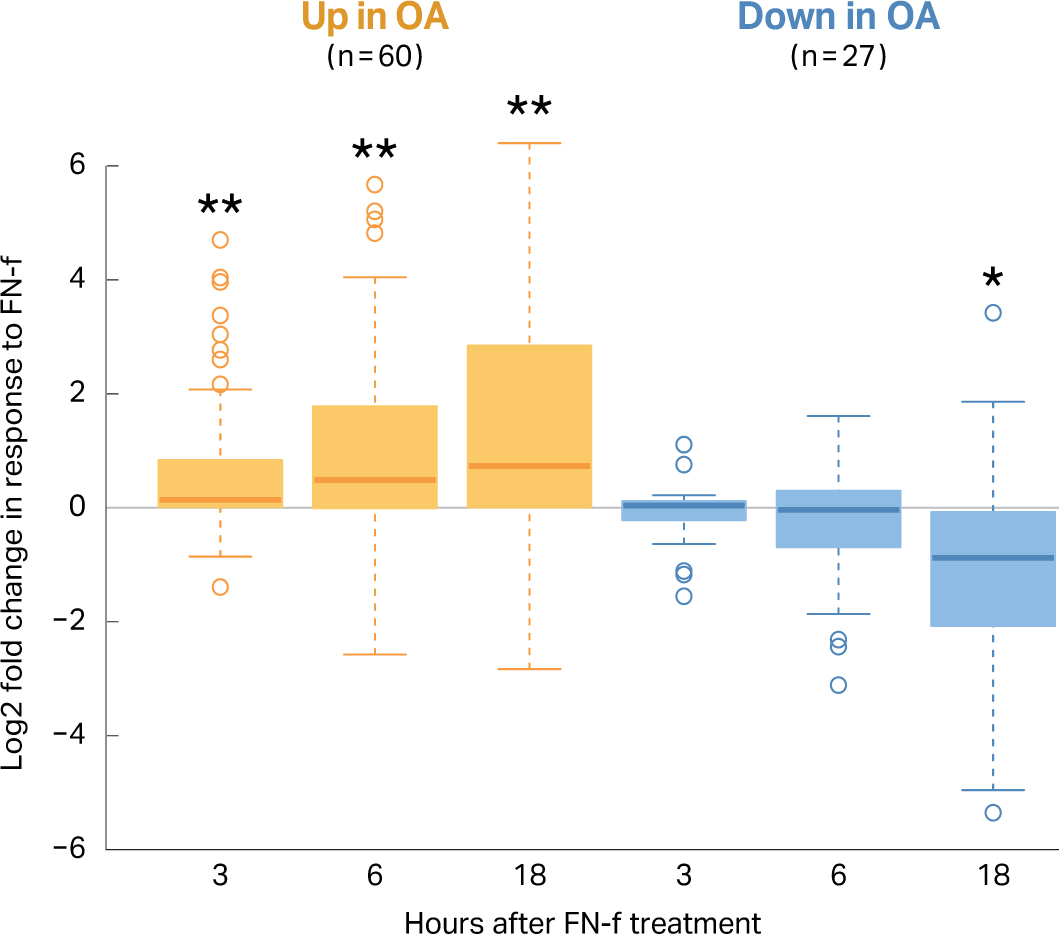
FN-f treatment induces changes similar to osteoarthritis. The previously published RAAK study compared osteoarthritis cartilage to preserved cartilage in the same joint, and of the differential genes reported, 60 were upregulated and 27 were downregulated with a high degree of change and significance (FDR p-value < 0.01, absolute fold-change > 1.5). The boxplot shows the log2 fold-change between FN-f treated and control samples at each time point for both the up- (yellow) and downregulated (blue) genes. These genes were ranked by fold-change and compared to a ranked list of fold-changes for genes outside of the OA set (Mann-Whitney p-values; ** =< 0.00001, * = 0.0004).

To further evaluate FN-f treatment as a model of the OA chondrocyte phenotype, we compared our differentially expressed genes to genes that have been implicated in OA by Genome Wide Association Studies (GWAS). Tachmazidou et al.^29^ recently identified 64 loci harboring OA-associated variants. These single-nucleotide polymorphisms (SNPs) represent regions with genotypes statistically associated with the OA phenotype, but most occur in non-coding regions of the genome rather than in gene bodies. This suggests that the SNPs likely impact regulatory regions, and the gene(s) they affect—which in turn promote the OA phenotype—could be hundreds of thousands of base pairs away. In order to identify genes that might be affected by these OA variants, we overlapped the locations of the SNPs with our differentially expressed genes and found that 72 of the genes identified here were within 500 Kb of a GWAS SNP. These genes are summarized in **Table 1** and include many genes with previously reported connections to OA, such as *NFKB1*^30^, *SOX7* ^31^, *IL11*^32^, *GDF5*^33,34^, *FGF18* ^35^, *TNFSF15*, and *NKX3-2* ^36^. These results reinforce the validity of this model for studying OA-related processes and could help identify the genes affected by OA-associated genetic variants.

**Table 1.**
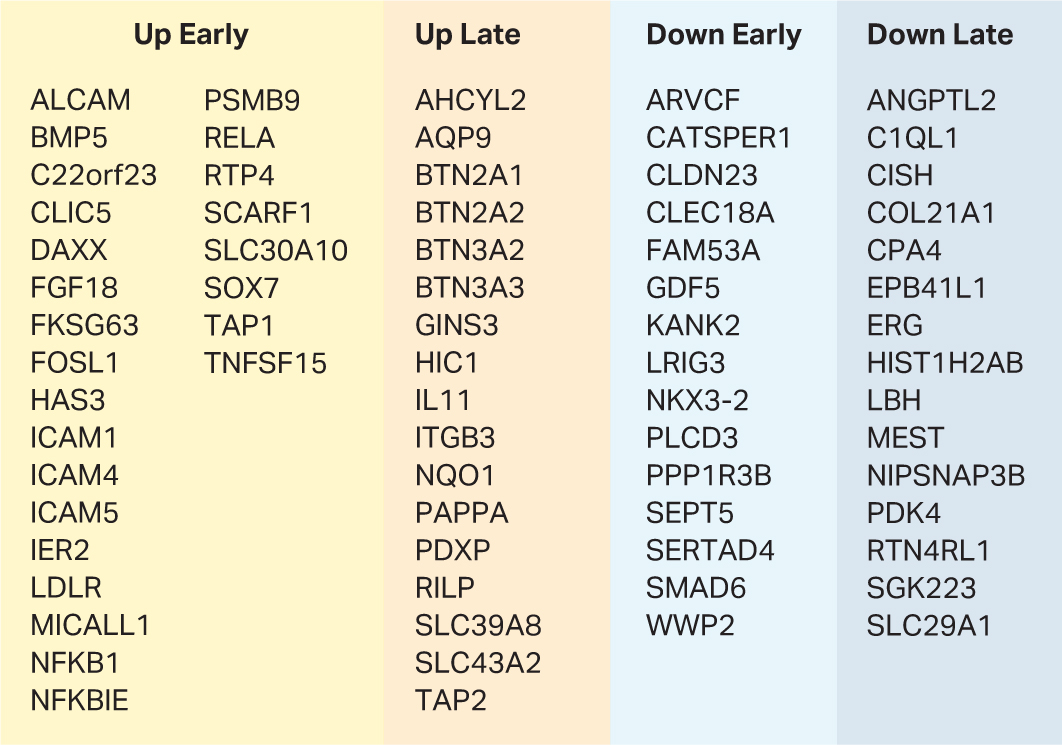
FN-f responsive genes near osteoarthritis GWAS loci. This table lists all differential genes that change in response to FN-f-treatment and that have a transcriptional start site within 500 Kb of a GWAS SNP as identified in Tachmazidou et al, 201929. For each gene, the response class is listed (as determined by k-means clustering of all differential genes)

### FN-f triggers both early- and late-response genes

To investigate the temporal patterns of transcriptional changes in response to FN-f, we performed k-means clustering of differentially expressed genes (**Fig 3**). This revealed four distinct temporal patterns. Early-response genes (both up- and downregulated) exhibited changes in expression as early as 3 hours but generally peaked at 6 hours of FN-f treatment. These early-response genes include many up-regulated genes that have been previously implicated in OA, including AP-1 components *FOS* and *JUN* ^24,37^, *NOD2*^38^, *RIPK2*^38^, the aggrecanase *ADAMTS4* ^24,39^, *CXCL1, CXCL2*, and *CXCL3*^16^, *LIF, TNF* ^40–42^, *TNFAIP6* and *TNFRSF11B*^19^, *PTGES*^19^, *IL6* and *IL8* ^39,41,43,44^. Among genes that decreased early was the transcription factor *SP7*, which is downregulated by TNF^45^. Late-response genes showed maximum absolute fold changes after 18 hours of FN-f treatment. Many of these genes have also been implicated in OA and/or matrix remodeling, including *MMP1, MMP10*, and *MMP13*^15,38,46^, *CD55, NGF* and *PAPPA*^19^, several interleukins such as *IL1B, IL11* and *IL17C* ^39,43,47^, the interleukin-receptor-associated kinase *IRAK3*, the IL1 receptor antagonist *IL1RN*, and collagens *COL13A1* and *COL7A1*^48,49^. The cartilage-specific integrin α10β1 (*ITGA10*) as well as integrin α11β1 (*ITGA11*) were both downregulated at late time points^50^. These results highlight the value of looking at FN-f response across a time course as it both reveals transient events not observed at every time point and provides insight into the temporal order and possibly even causal relationships between regulatory events.

**Figure 3.**
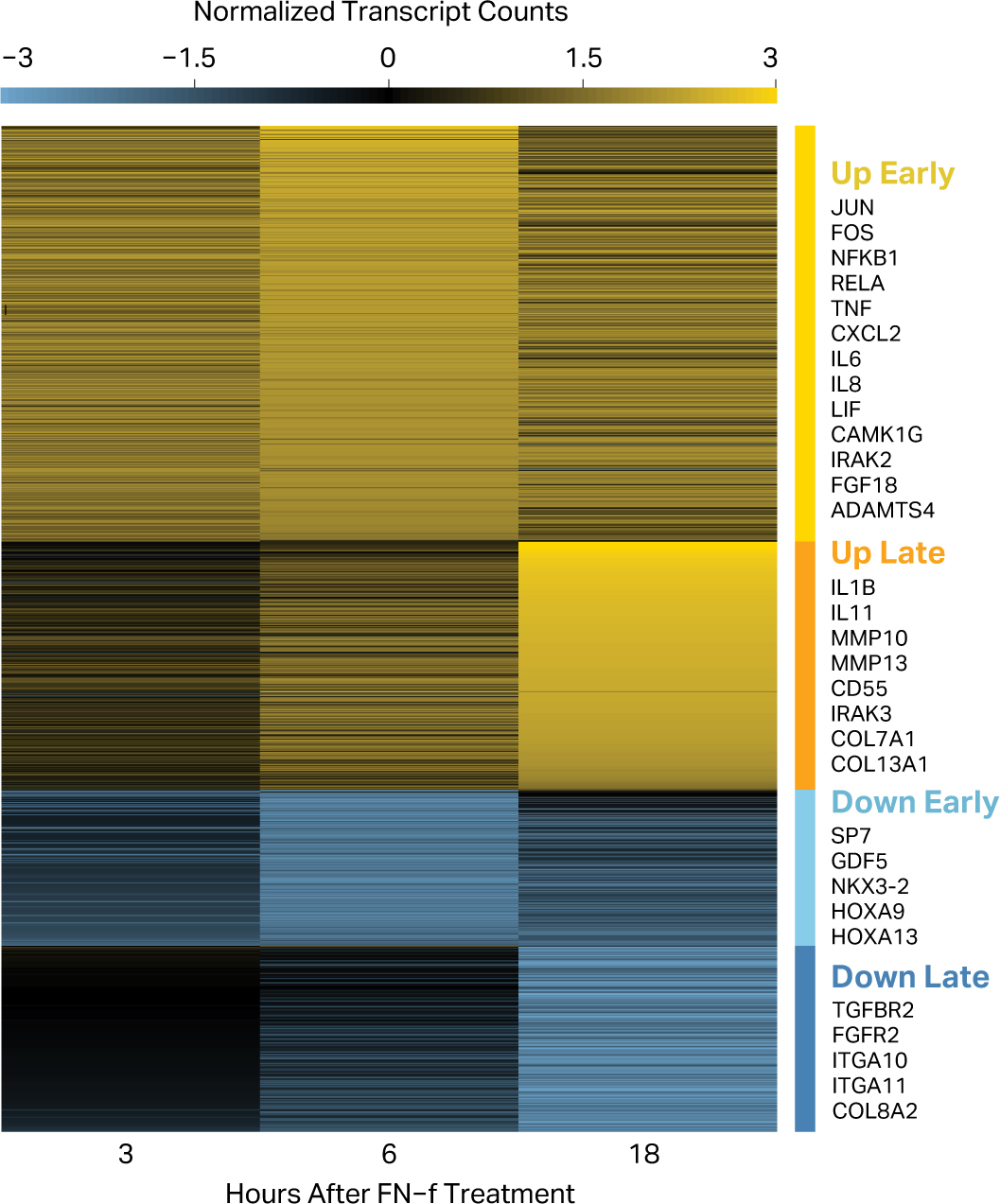
FN-f treatment regulates both early- and late-response genes. The genes that changed significantly in response to fibronectin fragment treatment were clustered according to their difference in z-score normalized counts between FN-f-treated and untreated samples at each time point. This separated the differential genes into four classes: “Up Early” (yellow; n=506), “Up Late” (orange; n=302), “Down Early” (light blue; n=190), and “Down Late” (dark blue; n=226). Selected genes are highlighted in each cluster.

We investigated whether the length of genes—and the corresponding time it would take to transcribe them—could account for the difference in response time. While the mean gene length for the late-response genes was statistically significantly longer than that of early-response genes (Mann-Whitney U test, p-value = 4e-6; **Fig S2**), the two distributions exhibited substantial overlap. Therefore, gene length is not likely to be the primary determinant of early vs late response.

### Fn-f induces transcription of proinflammatory genes and pathways

To understand the likely phenotypic impact of the changes induced by FN-f, we performed Gene Ontology (GO) enrichment analysis for the genes in each of the four clusters (**Fig 4A**). Both early- and late-response genes that were up-regulated in response to FN-f were strongly enriched for proinflammatory biological processes including “response to cytokine”, “inflammatory response”, and “immune system process”. Genes in these categories include NF-kB subunits, chemokine receptors, interleukins, MAP kinases, TNF ligands, and other cytokines. This is consistent with previous studies that have demonstrated that FN-f treatment stimulates a proinflammatory response via MAP kinases and NF-κB signaling^15,16,25,41,51^, as well as the established role of inflammation in the progression of OA^24,30,39,40,42,43,47,52,53^. Genes downregulated in response to FN-f were enriched for GO terms for development and morphogenesis and included HOX genes, *TGFBR2*, and *COL8A2*, among others.

**Figure 4.**
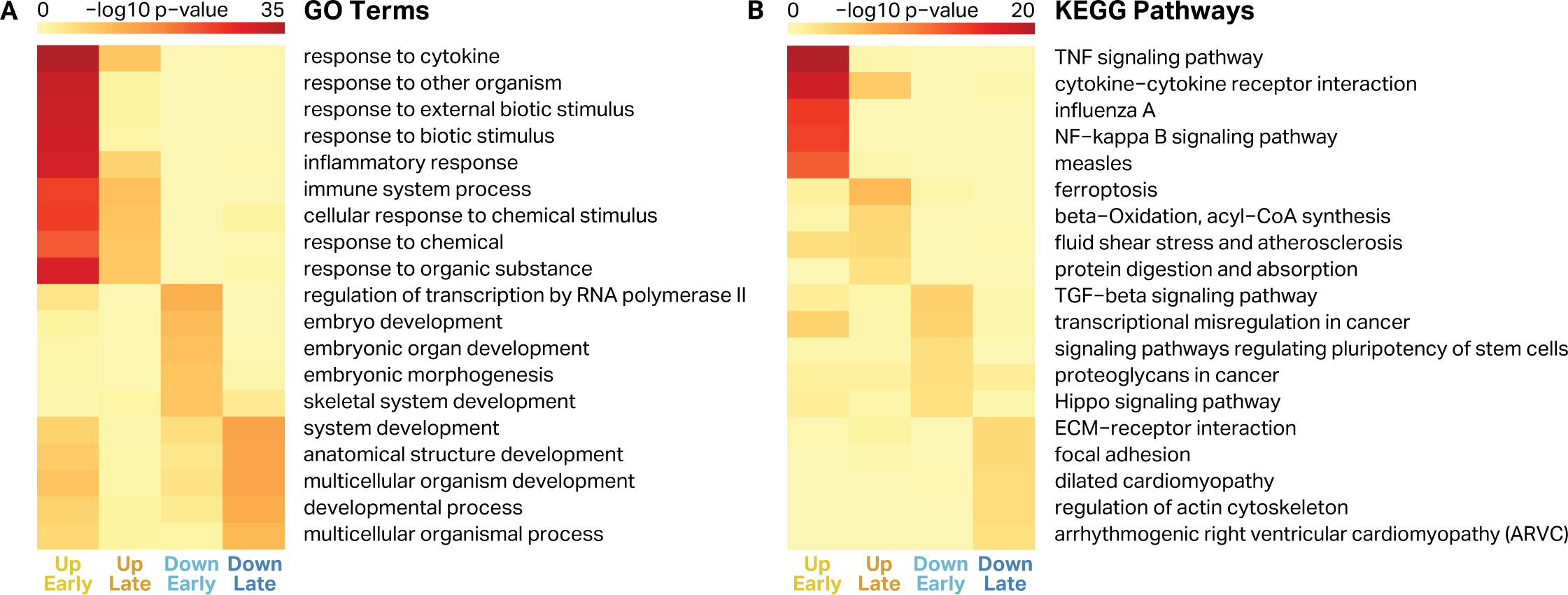
Gene ontology terms and KEGG pathways enriched in response to FN-f. For every cluster of differential genes, the top five GO terms **(A)** and KEGG pathways **(B)** were selected based on enrichment p-value. Heatmap color represents the -log10 of the enrichment p-value for each term or pathway for genes in each cluster (up early, up late, down early, and down late).

To determine pathways that were affected by FN-f treatment, we identified Kyoto Encyclopedia of Genes and Genomes (KEGG) pathways that were enriched in each of the four gene sets (**Fig 4B**). Early-response upregulated genes were enriched for TNF and NF-κB pathways, both of which have been implicated in OA and even targeted for therapeutic OA treatments^30,42,54^. Intriguingly, late-response upregulated genes were most strongly enriched for the ferroptosis pathway, a form of programmed cell death dependent on iron and accumulation of lipid peroxides induced by reactive oxygen species, which have been implicated in OA^55^. The late-response upregulated genes involved in ferroptosis included *ACSL1, GCLM, SLC39A8, HMOX1, ACSL5, FTH1, SLC7A11*, and *ACSL4*. The downregulated genes were enriched for the TGFβ and Hippo signaling pathway as well as ECM-receptor and focal adhesion genes.

To visualize how specific kinases were regulated in response to FN-f, we generated kinome tree maps using the human kinase visualization tool Coral (**Fig 5**). Kinase branches and nodes were colored to indicate time course clusters. This analysis identified many kinases with suspected roles in OA progression, including the p38 pathway member *MAP2K3*, which was upregulated 9.6-fold in response to FN-f^41,56^. In contrast, we found that *TGFBR2* was downregulated 3.5-fold in response to FN-f, which is consistent with recent findings in which decreased *TGFBR2* was correlated with increased OA severity in mice^57^. These studies also uncovered a number of kinases that have not been previously implicated in OA or chondrocyte dysfunction. For example, *DYRK3*, which was upregulated 3.7-fold in response to FN-f, is relatively poorly annotated and has thus been characterized as part of the “dark kinome”. *DYRK3* and other understudied kinases identified in this study provide novel targets for further study with regard to their involvement in OA.

**Figure 5.**
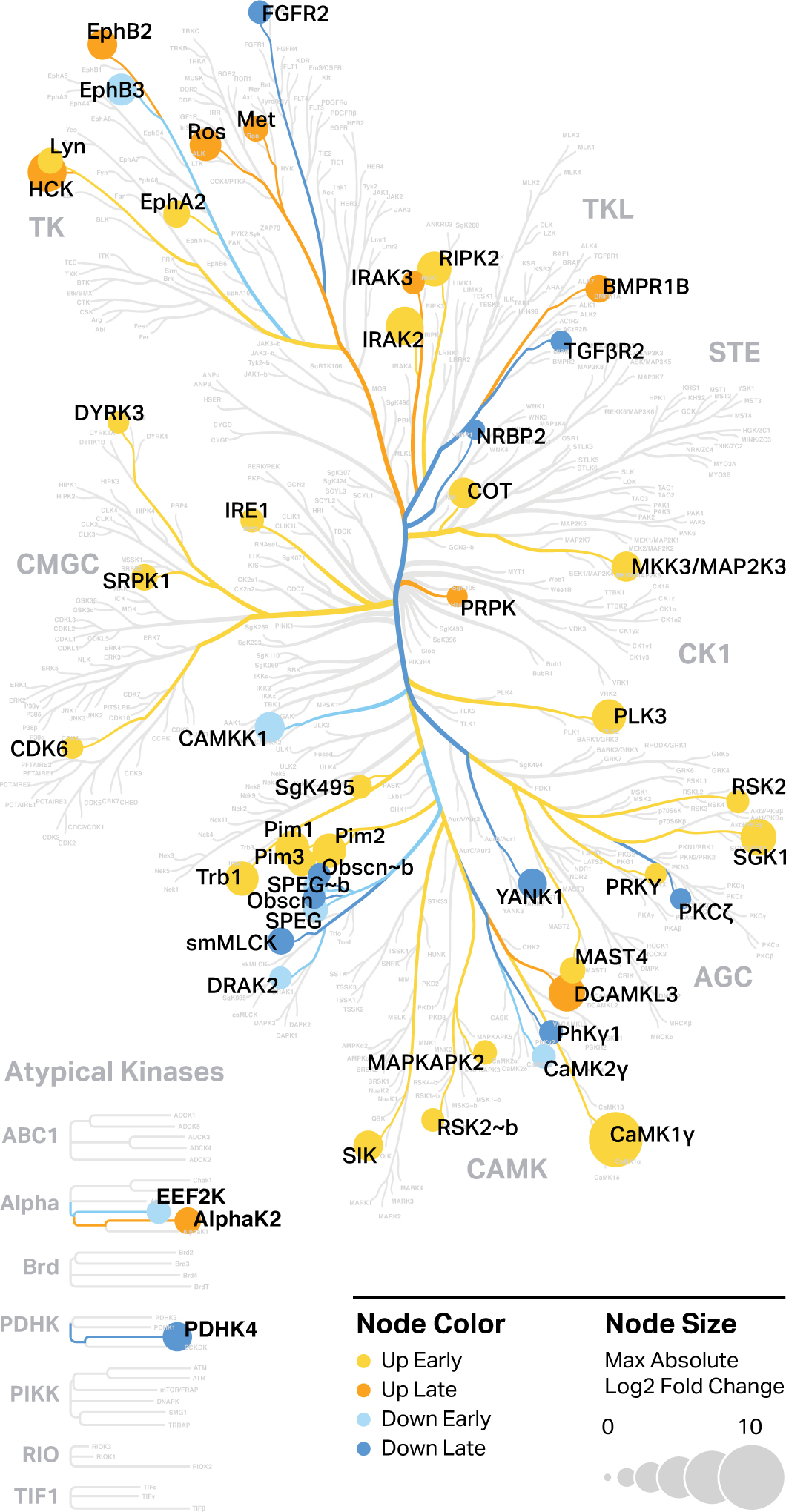
Protein kinases transcriptionally regulated by FN-f. Protein kinases that are differentially expressed in response to FN-f treatment were highlighted in a human kinome map generated by Coral. Color represents the temporal response class of the genes as determined by k-means clustering, while node size represents the maximum log2 fold change between time-matched FN-f-treated and control samples among the three time points.

### Transcriptional drivers of FN-f response include NF-κB & AP-1

To determine the transcription factors that were responsible for the global transcriptional changes induced by FN-f treatment, we used the HOMER software suite to identify transcription factor motifs that were enriched in the promoters of genes for each of the four clusters (**Fig 6**). Upregulated early-response genes exhibited a strong enrichment for motifs associated with multiple members of the NF-κB complex, including REL, RELB, TF65, and NFKB2. In contrast, upregulated late-response gene promoters were enriched for motifs of Jun-related proteins, including JUNB, NF2L2, and NFE2. JUNB is a member of the Activator Protein 1 (AP-1) which has been previously shown to contribute to OA and cartilage matrix degradation^24^. These findings are consistent with the fact that AP-1 and NF-κB subunits are also upregulated in response to FN-f.

**Figure 6.**
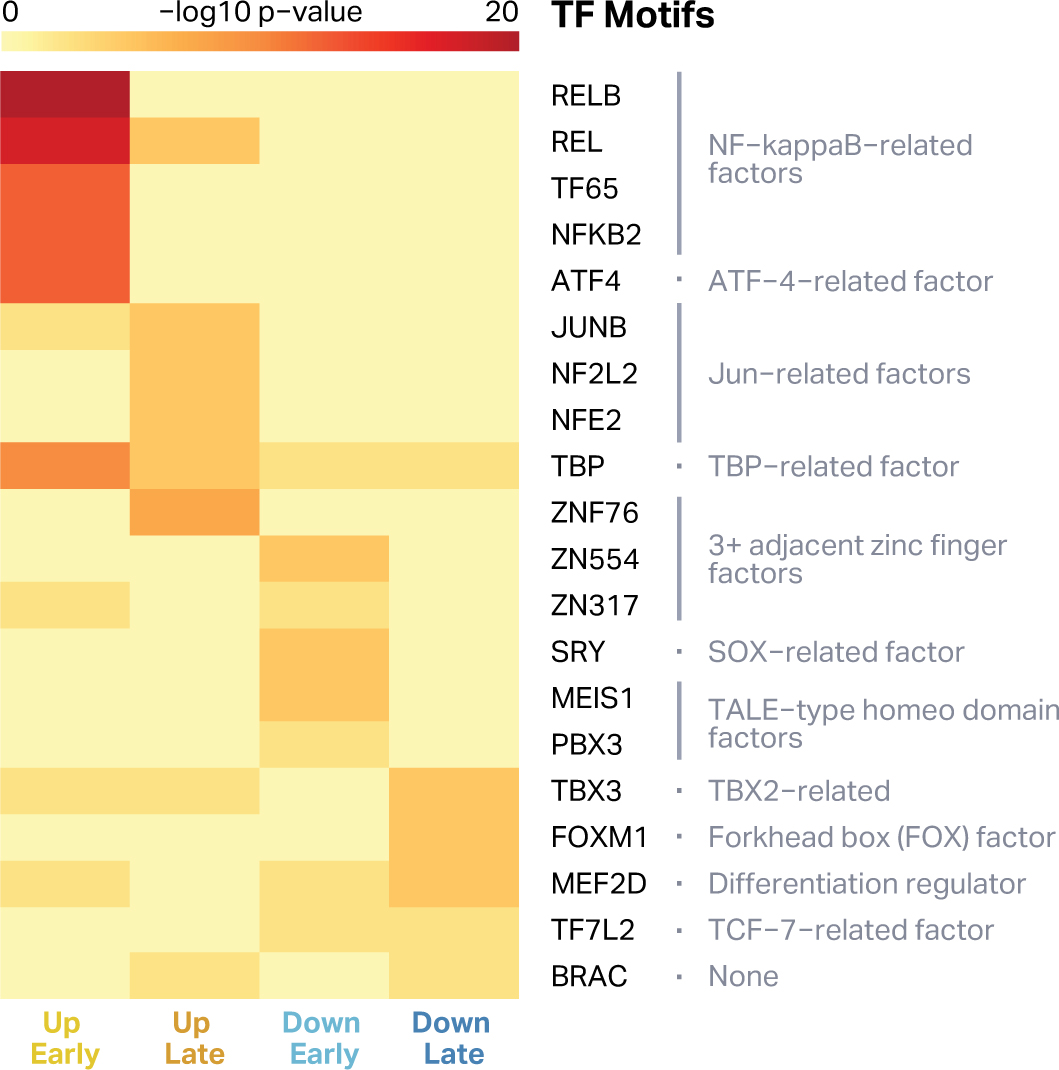
Transcription factor motifs enriched at FN-f early- and late-response genes. The promoters of genes in each cluster were analyzed for enriched transcription factor motifs from the HOCOMO-CO database, and the top five from each cluster (based on enrichment p-value) are plotted in the heatmap. Transcription factor families from TFClass are listed in grey on the right20. Each temporal class of genes has unique transcription factor motifs enriched. The promoters of genes upregulated early are enriched for the motifs bound by various NFkB members, while the promoters of genes upregulated late are enriched for factors related to the AP-1 component Jun.

This analysis also revealed the enrichment of several unannotated motifs in the promoters of upregulated genes (**Fig S3**). The presence of these *de novo* motifs may suggest that transcription factors with currently uncharacterized motifs also play a role in FN-f response. Further studies investigating the proteomic and phosphoproteomic landscape of chondrocytes responding to FN-f treatment may help in the further characterization of this model system and identify other key regulators in this response, which could also play a role in OA progression.

## Discussion

Developing cell culture systems that model key aspects of diseases can be incredibly valuable for deciphering mechanisms and testing therapeutic interventions, particularly when high-throughput screens are necessary. Using a fragment of the matrix protein fibronectin, we investigated a human cell-culture model of chondrocyte response to cartilage matrix breakdown, a key trigger of OA. Our transcriptome-wide analysis confirmed a similarity between this system and changes observed in OA tissue, suggesting that this is a powerful system with which to study the OA chondrocyte phenotype. We confirmed that many OA-responsive genes also change significantly in normal chondrocytes in response to FN-f. In addition, we classified over a thousand FN-f-responsive genes by their direction and timing of regulation and confirmed that many genes and pathways upregulated in response to FN-f have previously been characterized as a part of the OA phenotype. This includes inflammatory cytokines and chemokines such as *IL1B* and CXC ligands, matrix-degrading proteinases such as *MMP13*, and members of the NF-κB signaling pathway.

The presence of early- and late-response gene clusters in this system is reminiscent of primary and secondary responses observed in other systems, such as the inflammatory response in immune cells^58–63^ and the growth factor response promoting differentiation and proliferation^64^. Variation in response times can stem from differences in genomic and regulatory features, including the degree to which products are regulated at transcriptional, post-transcriptional and translational levels, the baseline differences in RNA polymerase II occupancy at transcription start sites, and the dependency of target gene regulation on epigenetic modifications^64^. Further exploration of these and other regulatory mechanisms in this system may provide a deeper understanding of how chondrocytes are phenotypically altered in OA.

This study also revealed many genes and transcription factors that have not been previously associated with OA, suggesting new lines of interest for further investigation. One example is the finding of over-representation of the ferroptosis pathway in the upregulated late-response genes. Ferroptosis is a relatively recently described mechanism of cell death that involves iron and excessive levels of lipid peroxides generated by oxidation of lipids^55^. Ferroptosis can result from disturbances in the glutathione-dependent anti-oxidant system, release of excessive reactive oxygen species (ROS) from the mitochondria, and oxidation of lipids by lipoxygenases and cyclooxygenases^65^. Previous studies have demonstrated that FN-f treatment of chondrocytes generates ROS that regulate signaling involved in MMP expression^18^. Although ferroptosis per se has not been described in OA cartilage, studies have demonstrated lipid peroxidation^66^, glutathione oxidation^67^, mitochondrial dysfunction^68^, and increased activity of lipoxygenases and cyclooxygenases^69^, indicating ferroptosis could contribute to chondrocyte death in OA cartilage. This finding may also be relevant to osteoarthritis associated with hemochromatosis where excessive iron is present^70^.

The intersection between FN-f-responsive genes and OA GWAS loci provides a subset of genes that could be affected by OA-associated genomic variants, offering potential targets for follow-up studies. In addition to the NF-κB family, which has been considered as an OA target for quite some time, *FGF18* was a FN-f-responsive gene (up early) present in the GWAS dataset. Unlike NF-κB, *FGF18*, which is an anabolic growth factor, is in clinical trials for knee OA as an intra-articular agent that may promote cartilage growth^71^. Additional growth factors present in both datasets were the BMP family members *GDF5* (down early) and *BMP5* (up early) as well as the BMP signaling protein *SMAD6* (down early). Consistent with the FN-f-induced chondrocyte phenotype, allelic variation in the *GDF5* gene has been associated with reduced expression^72^. *BMP5* is a regulator of bone and cartilage formation during development, but its role in OA is not clear^73^. *SMAD6* is an inhibitor of *SMAD1/5*, and its overexpression in mice was associated with a reduction in osteophyte formation, suggesting that decreased *SMAD6* expression could be detrimental in OA^74,75^.

While FN-f treatment of *ex vivo* chondrocytes represents a powerful tool to understand some of the events that promote OA, it does not recapitulate all aspects of OA, nor does it serve to replace animal models nor analysis of human tissue. Osteoarthritis is a complex disease involving multiple tissues and arises due to both genetic and environmental factors. A more complete mechanistic understanding of OA therefore requires orthogonal approaches with offsetting advantages and limitations. This *ex vivo* FN-f treatment model does, however, fill a valuable gap and provide a flexible and manipulatable system with which to understand the behavior of chondrocytes in both healthy and disease conditions. By combining this system with recent advances in genomics and genome editing (including the ability to edit primary human chondrocytes^76^), this FN-f model offers incredible promise for study of OA.

## Acknowledgements

We would like to thank the Gift of Hope Organ and Tissue Donor Network, and the donor families, for providing normal donor tissue. We would like to thank Dr. Arkady Margulis for donor tissue procurement and Mrs. Arnavaz Hakimiyan for technical assistance. We would also like to thank Erika Deoudes for figure and preprint design.

## Author Contributions

K.S.M.R. and C.K. carried out the analysis and interpretation of the data.

K.S.M.R. and V.U. collected and assembled the data.

K.S.M.R., D.H.P., and R.F.L. drafted the manuscript.

C.K. and V.U. revised the article for important intellectual content.

R.F.L. and D.H.P. obtained funding for, conceived, and designed the experiments.

All authors provided final approval of the article prior to submission.

## Supplemental Figures

**Figure S1.**
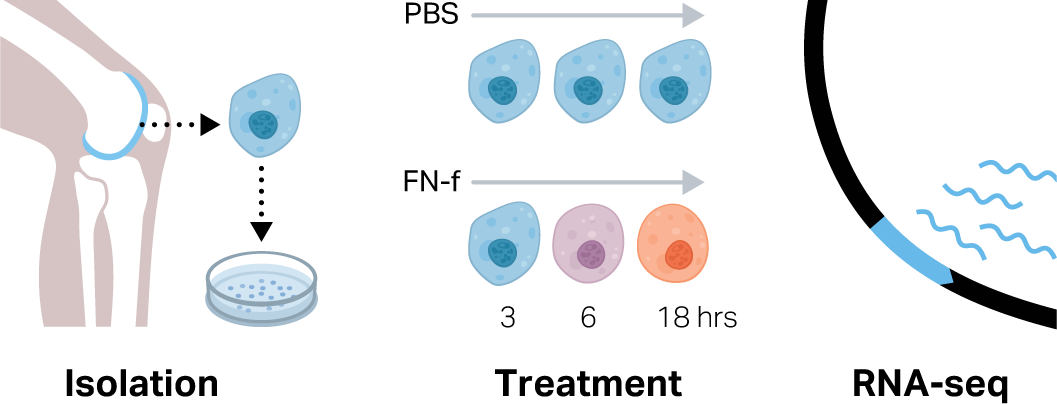
Experimental design of fibronectin fragment (FN-f) treatment. Primary articular chondrocytes were isolated from normal human femoral cartilage obtained from three tissue donors aged 50-61 years, each without osteoarthritis. Six samples from each donor were cultured and treated with either 1 µM fibronectin fragments (FN-f) or PBS. After 3, 6, or 18 hours, RNA was extracted and used to create RNA-seq libraries. This design allowed for the comparison of gene expression in FN-f-treated samples compared to time-matched untreated controls, accounting for changes occurring as a result of isolation, time in culture, and donor genotype.

**Figure S2.**
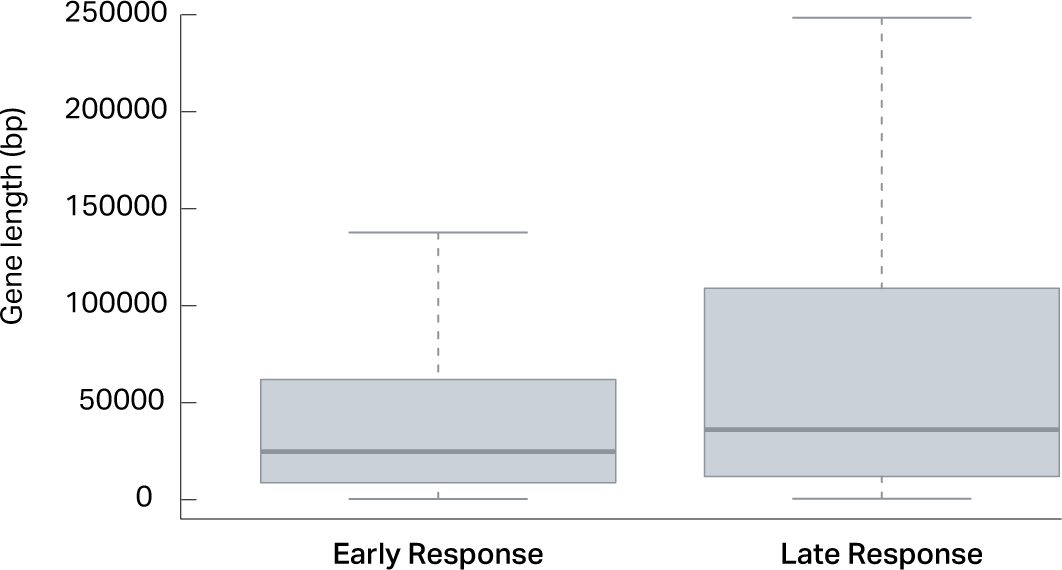
Gene length distributions among genes up- and down-regulated by FN-f. Boxplots depicting the lengths of early-response genes (both up and down, n=696) and late-response genes (both up and down, n=528) reveal that late-response genes are statistically significantly longer (Mann-Whitney U test, p-value <0.0001). Outliers are excluded from this plot.

**Figure S3.**
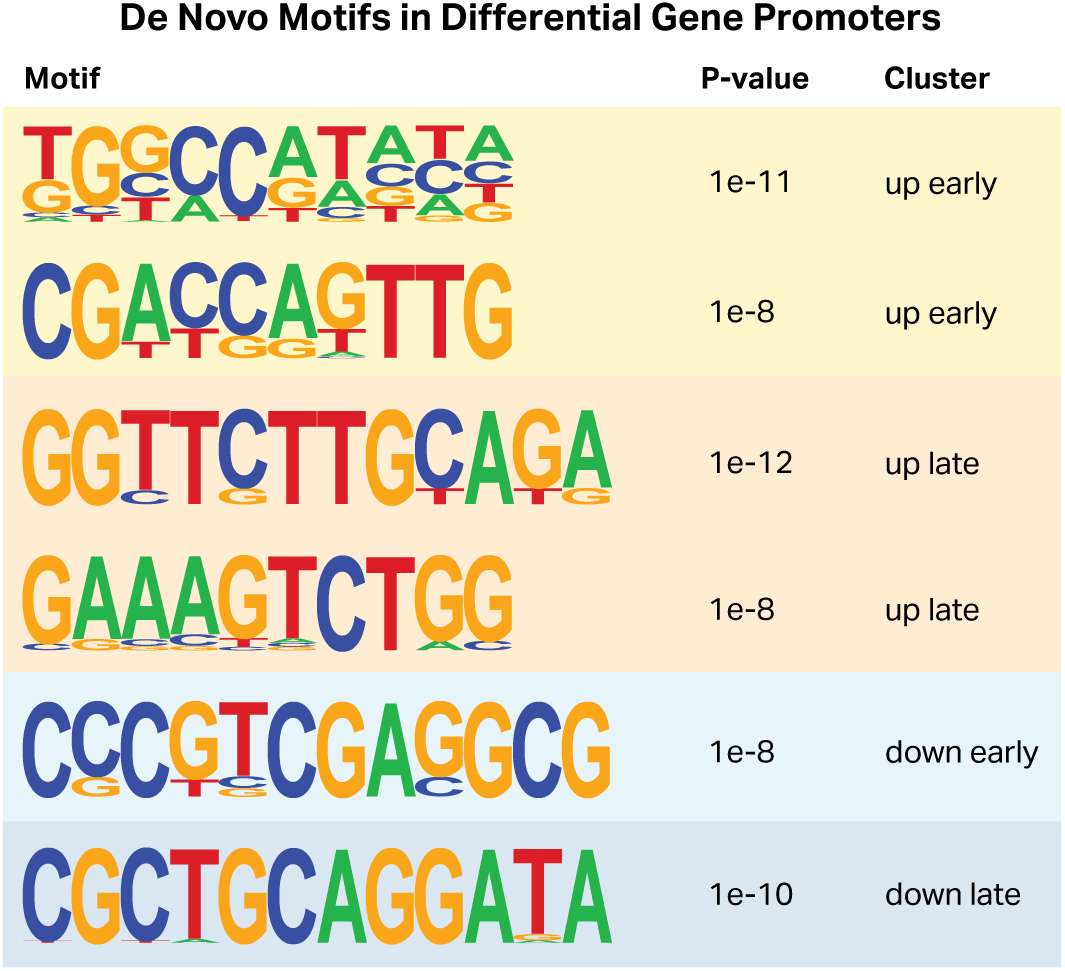
Unannotated *de novo* motifs identified in differential gene promoters. When the promoters of differential genes among each cluster were analyzed for enriched transcription factor motifs, several had no clear match to known motifs (motif match score < 0.6). These unannotated *de novo* motifs represent potential binding sites of transcription factors that have yet to be characterized. For each motif, the motif logo, p-value of enrichment, and cluster in which it was enriched are listed.

**Table S1. Differential genes in response to FN-f treatment**

This table lists all differential genes (n=1,224) that change in response to FN-f treatment. For each gene, the response class is listed (as determined by k-means clustering of all differential genes), as well as the p-value (Wald test), log2 fold-change, and z-score normalized counts at each time point, and the raw counts from each sample.

**Table S2. Osteoarthritis genes in response to FN-f treatment**

This table includes all OA-responsive genes from the RAAK study with a p-value less than 0.01 and an absolute fold-change greater than 1.5 (n=87), and details how they change in response to FN-f treatment. For each gene, the response class is listed (as determined by k-means clustering of all differential genes), as well as the p-value (Wald test), log2 fold-change, and z-score normalized counts at each time point, the raw counts from each sample, and the original p-value and fold-change from the RAAK study [19].

